# Characterizing Functional Clusters of V4 Neurons in Digital Twins

**DOI:** 10.64898/2026.07.25.740202

**Authors:** Rendong Tang, Qiongyi Zhou, Wenghao Zhao, Zhuoyue Dai, Rui Zhang, Jiayu Wang, Gongting Wang, Changde Du, Huiguang He, Haidong D. Lu

## Abstract

Neurons in primate visual cortical area V4 display tuning for multiple visual features, including color, shape, texture, and depth. Whether and how these neurons are organized into functional architectures remains largely unknown. Using two-photon calcium imaging in anesthetized macaques, we recorded responses of hundreds of V4 neurons to natural images and used these data to train deep convolutional neural network models (digital twins), obtaining synthetic images (SI) that maximized neuronal responses. Based on their SIs, these neurons clustered into classes that spatially matched the orientation, color, and curvature maps from intrinsic signal imaging. Furthermore, lesion study in digital twins revealed different integration rules for different neuron classes. Thus, digital twins of V4 neurons can be applied as a promising tool to characterize neurons’ fundamental features, which underlie the functional clustering in this area.

**Highlights:**

- Neurons in V4 are examined with two-photon calcium imaging and DCNN modeling (digital twins)
- Synthetic images derived from neurons’ digital twins reveal distinct neuron groups
- These groups match functional types defined by intrinsic signal optical imaging
- Lesion study in digital twins demonstrates neural mechanisms underlying feature tuning

## Introduction

The primate V4 is an intermediate visual area responsible for processing color, medium-complexity shapes, and attentional modulation (Roe et al. 2010; Pasupathy et al. 2020). Unlike V1 and V2, V4 lacks distinct cytochrome oxidase (CO) compartments. Whether V4 possesses functional architectures that similar to the columns in V1/V2 has been a subject of debate. On the one hand, imaging studies in V4 have revealed clear functional maps for color, orientation, and curvature (Ghose & Ts’o 1997; Conway et al. 2007; Tanigawa et al. 2010; Li et al. 2014; Tang et al. 2020). On the other hand, single-unit electrophysiological studies have suggested that clustering in V4 either does not exist (Schein et al. 1982) or is sparse (Kotake et al. 2009; Namima et al. 2025), since neighboring neurons often exhibit significant differences in their selectivity (e.g., Namima et al. 2025). Such a discrepancy may due to multiple reasons. First, neurons in a region are only sparsely sampled in electrophysiological studies and it is difficult to estimate the response properties of population of neurons within an area. Second, and perhaps more importantly, traditional visual stimuli are limited in characterizing complex V4 neurons, making it difficult to judge the similarity of neighboring cells. A single V4 neuron is typically tuned to multiple stimulus dimensions (e.g., orientation, color, disparity, texture), and the neuron may respond best to a particular combination of these features. Within limited recording time, however, traditional recordings can only examine a limited part of the parameter space.

Recently, digital twins of visual neurons have been developed to better characterize neurons’ tuning features (Bashivan et al. 2019; Walker et al. 2019). Researchers first record cell’s responses to a large set of natural images, then train a neuron model built upon a deep convolutional neural network (DCNN). The resulting single-cell model (digital twin) can accurately predict the cell’s responses to new stimuli. Cell-specific synthetic images (SIs; Bashivan et al. 2019), or most exciting inputs (MEI, Walker et al. 2019) can be derived from the trained models. These SIs have complex patterns that are very different from conventional stimuli. Many cells respond more strongly to the SIs than to the natural image tested. Thus, SIs can be considered a more accurate description of a cell’s receptive field properties than traditional methods. Using this technique, one study showed that neighboring V4 cells have similar SIs, forming neuronal clusters (Willeke et al. 2026). However, studies on this topic were mainly based on electrophysiology, and had limited spatial information about cell distributions. To overcome this limitation, two-photon imaging is an ideal method for directly studying the spatial topology of cell populations. In this study, we combined two-photon calcium imaging and DCNN modeling to study areas V1 and V4 in anesthetized monkeys. Furthermore, we used intrinsic signal optical imaging (ISOI) to obtain functional maps on the V4 surface, aiming to link single-cell selectivity with traditional functional maps through two modalities of imaging on the same cortex.

Another important question we aim to address concerns how multi-selective V4 neurons process mixed features, i.e., the degree of feature binding at the single-cell level. For example, regarding the relationship between color and shape, some studies have found they are independent (Schein et al. 1982; Bushnell & Pasupathy 2012), while others have found they have interactions (McClurkin & Optican 1996; Zeki 1983). Recent DCNN studies, using grayscale natural images, have not addressed this issue (Bashivan et al. 2019; Walker et al. 2019). In the present study, we used color images to test the relationship between color and shape.

## Results

### Two-photon imaging captures single-neuron responses across V4 functional domains

Data in this study are collected from three imaging chambers in three macaque monkeys (Case 1-3, Figure S1). Animal preparation and imaging procedures were similar to those described in our previous studies (Tang et al. 2020; Zhang et al. 2025; Wang et al. 2025). We first performed intrinsic signal optical imaging (ISOI) to identify three typical types of functional domains in V4 (orientation, color, curvature; Figure 1A-E). Then, we injected AAV virus (AAV9.Syn.GCaMP6S) into different functional domains. Injection depth was 500 μm, and injection volume was 500 nL per site. Six weeks later, a 13 mm-diameter-imaging chamber was installed.

**Figure 1.**
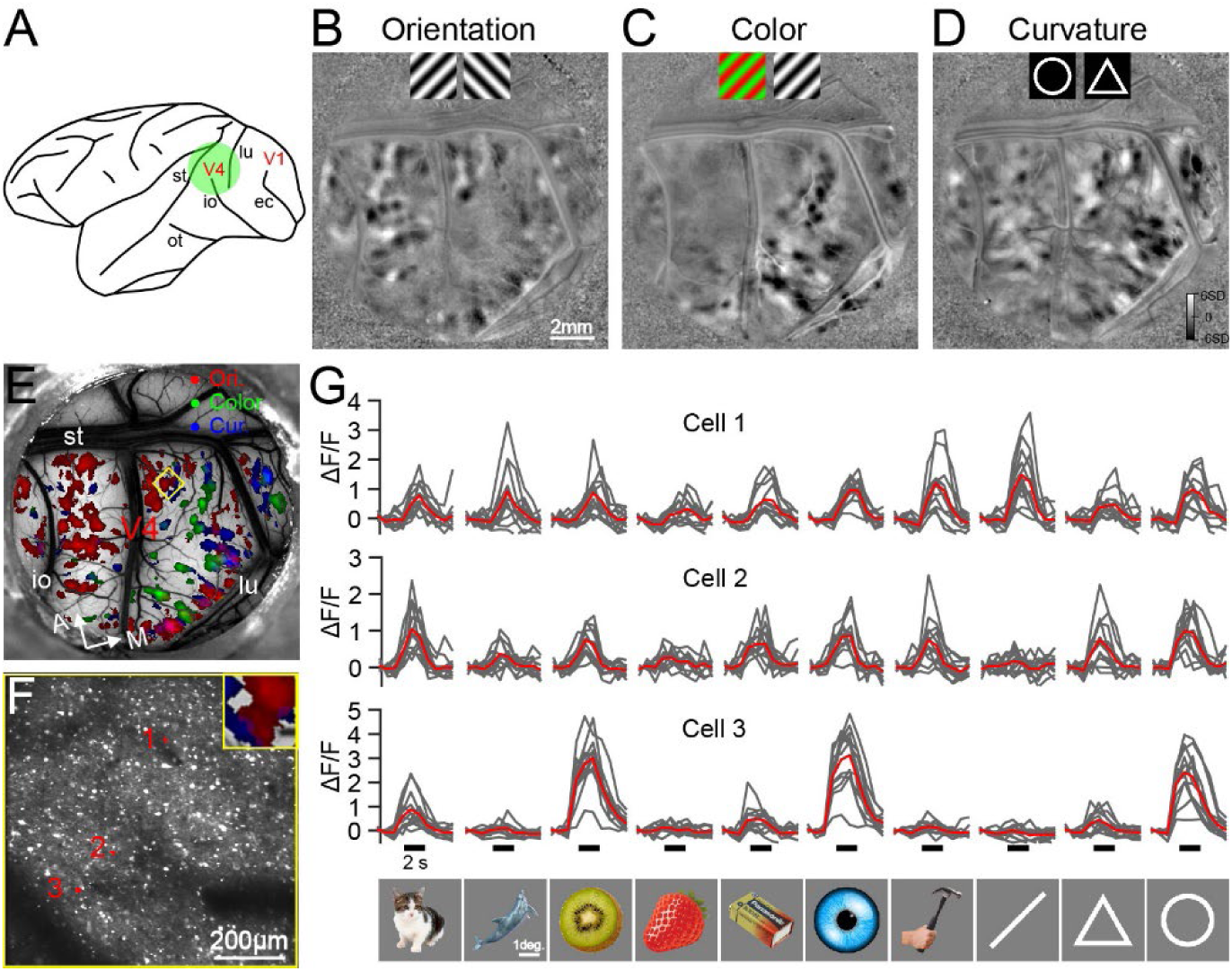
ISOI functional maps in V4 and neuronal calcium responses to visual images. **A**: An illustration of a monkey brain and a chamber located over area V4. lu, lunate; st, superior temporal; ec, external calcarine; io, inferior occipital; ot, occipitotemporal. **B-D**: Functional maps obtained with ISOI. **B**: Orientation map obtained by comparing responses to two gratings of orthogonal orientations. Scale bar applies to B-E. **C**: Color map obtained by comparing responses to red-green gratings and black-white gratings. **D**: Curvature map obtained by comparing responses to circles and triangles. **E**: Three types of V4 functional domains derived from ISOI results (B-D): red for orientation domain, green for color domain, blue for curvature domain. Yellow box indicates a two-photon imaging site, shown in panel F. **F**: Two-photon fluorescence image of the V4 site corresponding to the yellow box in E, imaged from depth 225 μm. The far-left side of this site is in a curvature domain, and the mid-right part is in an orientation domain (see inset). A total of 118 cells were identified from this imaging region. Responses of three example cells (red dots) are shown in G. **G**: Calcium fluorescence responses of three example V4 cells to 10 typical stimuli. Black short lines indicate stimulus presentation time (2 sec). Each stimulus was repeated 12 times (gray curves), with the red curve showing the average response.

In the subsequent two-photon calcium imaging, we used a 16X objective, with imaging frequency of 1.3 Hz in Galvo mode. Imaging area covered 0.83 × 0.83 mm², and each frame consisted of 512 × 512 pixels. We performed two-photon imaging at 8 sites across the 3 chambers (3 V1 sites and 5 V4 sites; Figure S1), with imaging depths ranging from 155–295 μm. Each field of view (FOV) yielded on average ∼100 active cells (Figure 1F). We occluded one eye, stimulated only the other eye, and determined the location of the neuronal population receptive field for the imaged area. For the corresponding cortical visual field, V1 population receptive field sizes were typically ∼1.3 degrees, and V4 receptive fields were 4.5–7.5 degrees. Visual stimuli comprised 800 images of natural objects, and for 5 V4 sites, an additional 50 images of geometric figures (e.g., bottom of Figure 1G). Stimulus size was 5 degrees for V1, and 80% of the population RF size for V4. Each stimulus was presented for 2 seconds, with an inter-stimulus interval of 3 seconds. Stimuli were displayed in a random order and each was repeated at least 10 times.

##Figure 1G shows calcium fluorescence responses of three example V4 cells to 10 stimuli; these cells were from different functional domains within the same V4 site (Figure 1F). Cell 1 located in an orientation domain, responded strongly to a 45° oriented stimulus. Cell 2, in contrast, responded to many stimuli but weakly to the 45° oriented stimulus. Cell 3, located in a curvature domain, responded strongly to several circular stimuli (except for the strawberry) but weakly to oriented or angled images (e.g., triangles). Overall, cells exhibit some distinct feature preferences (e.g., orientation and curvature), yet also share similarities, as many stimuli evoke responses to certain extents in all cells. The preferred stimulus features of these cells are not so obvious from these responses.

### Cell-specific synthetic images reveal distinct feature preferences of V4 neurons

In order to characterize the response preferences of cells, we built single-neuron DCNN models following the published work (Bashivan et al. 2019; Figure 2A). Briefly, the first three convolutional layers (L1-L3) of a pretrained AlexNet were used to extract features from the stimulus images; then followed by a “what” layer consisting of the channel weights of the final convolutional layer and a “where” layer corresponding to the cell’s spatial weights of its receptive field (Wang et al., 2020). We compared the resulting output (predicted ΔF/F) with the cell’s actual ΔF/F response and used error backpropagation to adjust the network connection weights (see Methods). We tried using different numbers (1–3) of convolutional layers to build the networks. For V1 neurons, using only L1 or L1+L2 yielded the best fit; for the V4 neurons, using all 3 layers (L1-L3) yielded the best fit (see Methods). Figure 2B shows the comparison between model-predicted responses and actual responses for one V1 cell and one V4 cell. The trained models simulate cells’ responses well (left panels) and also predict their responses to new visual stimuli well (right panels). We used EVE (Explainable Variance Explained) to measure the goodness-of-fit of the models.

**Figure 2.**
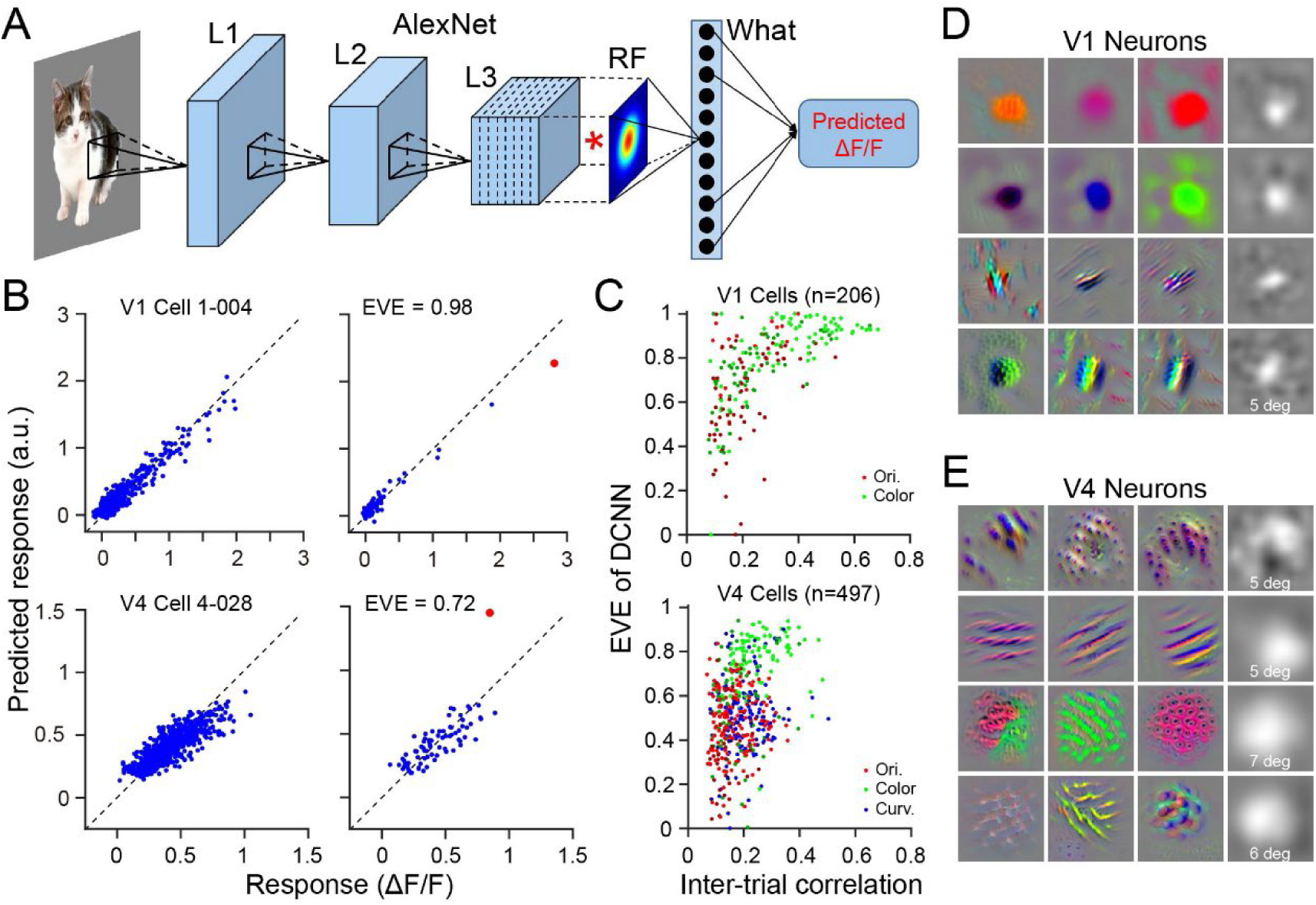
DCNN models for V1 and V4 neurons. **A**: Model architecture. Similar to the published work (Bashivan et al. 2019), the single-cell model is based on a pretrained AlexNet, with “where” (RF) and “what” layers added next to the convolutional layers L1-L3. The diagram shows the model for V4 neuron, as the V1 model does not include L3. **B**: Comparison between actual responses (x-axis) and model-predicted responses (y-axis) for two example cells. Left column is for training set stimuli (720 stimuli) and right column is for the test set (80 stimuli). Dashed lines represent the diagonals. Red points indicate responses to the SI stimuli. **C**: Upper panel: Relationship between V1 neuronal response stability (average correlation coefficient across repeats; x-axis) and DCNN prediction accuracy (EVE; y-axis). Different cell types are shown in different colors. Note: cells with correlation values <0.075 were excluded from model fitting analysis (same below). Bottom panel: same for V4 cells. **D**: Columns 1–3 are SIs of 12 example V1 cells from color domains (rows 1-2) and orientation domains (row 3-4). Column 4 shows RF images of the neurons in column 3, i.e., the “where” layer weights in the DCNN model. Each image is 5 degrees in width. **E**: Same as D, but for 12 example V4 neurons: curvature cells (row 1), orientation cells (row 2), color cells (row 3), and cells from outside of the functional domains (row 4). Column 4 shows RF images of the neurons in column 3. Images sizes are 5 or 7 degrees apply to the same row.

Goodness-of-fit was related to the stability of the cell’s responses, particularly evident in V1 cells (Figure 2C, upper panel). Stability was measured as inter-trial correlation for stimulus set over different repeats. The overall goodness-of-fit for V4 was lower than for V1, indicating that V4 cells have higher response complexity and are more difficult to fit than V1 cells. Within V4, color cells had higher goodness-of-fit than orientation and curvature ones (Figure 2C, bottom panel). In total, 497 V4 cells and 206 V1 cells with inter-trial correlation coefficient higher than 0.075 were modeled and included in further analysis.

We further used the trained models to generate SIs for each cell (see Methods). With a random noise pattern as the initial stimulus input, we used gradient ascent to optimize the stimulus image in pixel space, with the goal of maximizing the model output, i.e., the cell’s predicted maximal response. The final optimized stimulus image is the cell’s SI. Figure 2D and 2E show SIs of 12 V1 cells and 12 V4 cells, respectively. The receptive fields of V1 cells were relatively small (mean size 1.4 deg), and the spatial structure within the receptive fields was relatively simple. Notably, the receptive fields of color cells appeared as relatively uniform circular spots. Unlike V1 cells, V4 cells had relatively large receptive fields (mean size 3.9 deg), and the spatial patterns of their SIs were more complex. SIs of cells from different V4 functional domains (different rows in Figure 2E) showed substantial differences (see also Figure S2A).

Plotting the SIs at the cells’ cortical positions (Figure 3B) revealed that neighboring cells had similar SIs. Figure 3 shows an example site that located in a color domain (Figure 3A), comprising three subregions with different color preferences, within which cells preferred red, green, or blue (Figure 3B). Within these 200-500 μm subregions neurons prefer similar colors. Using different random initial images produced similar SIs (Figure 3C&D, Figure S2A). In the transition zones between these subregions, some cells preferred both colors (dual preference; top and bottom arrows in Figure 3B, and top and bottom rows in Figure 3D), while others exhibited mixed single-color preferences, such as the brown SI at the red-green transition border (middle arrow in Figure 3B, middle row in Figure 3D). These may reflect two different modes of combining color inputs (Nigam & Dragoi 2021). In orientation and curvature domains, neurons also have common characteristic SI patterns for orientation and curvature features (Figure S3).

**Figure 3.**
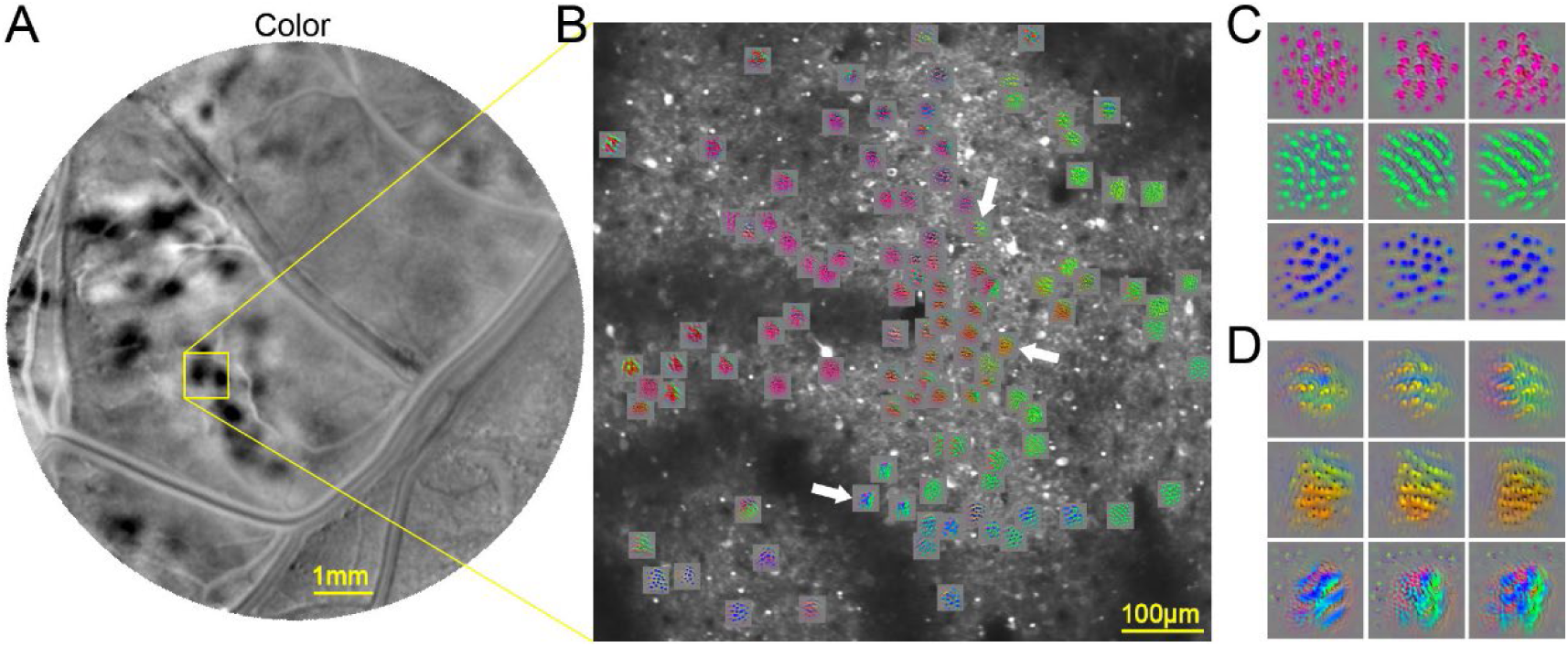
SIs of cells in a V4 color domain. **A**: Distribution of color domains within the imaging chamber of Case 3 (ISOI, also see Figure S1). **B**: Fluorescence image of a V4 color site and SIs of some cells. Cells’ SIs reveal subregions with different color preferences. Arrows indicate neurons located on boundaries between color subregions, where different color mixing patterns are observed. These neurons were shown in D. **C**: Three example cells (three rows) with different color preferences. Each show three SIs obtained from three different SI calculation sessions (three columns). **D**: Similar to B, showing SIs of three example cells (arrows marked in B). Unlike the cells in B, the preferences of these three cells are mixtures of two different colors.

### Unsupervised clustering of Sls recapitulates lSOl functional maps at single-cell resolution

Figure 4A shows more SI examples for V4 cells from three different sites: an orientation site (left), a color site (middle), and a curvature site (right). Within each site, the 12 SIs are similar (top 3 rows): the 12 cells in the orientation site share similar orientation preferences, and the 2D FFT frequency plots (row 5) of the representative cells (row 3) also shows similar spatial orientation features. Cells in the color site prefer similar colors, with a granular spatial structure but no prominent orientation features. Cells in the curvature site exhibit no clear orientation or color features but all show a certain curved structure. Row 4 in Figure 4A shows the RF images of the neurons in row 3, i.e. the weight map of the “where” layer in the model. The receptive fields of orientation and color cells are approximately circular, with stronger central weights and weaker peripheral weights. Curvature cells, in contrast, exhibit a ring-like pattern with weak center and strong periphery. Color cells have the largest receptive fields (scale bars are different).

**Figure 4.**
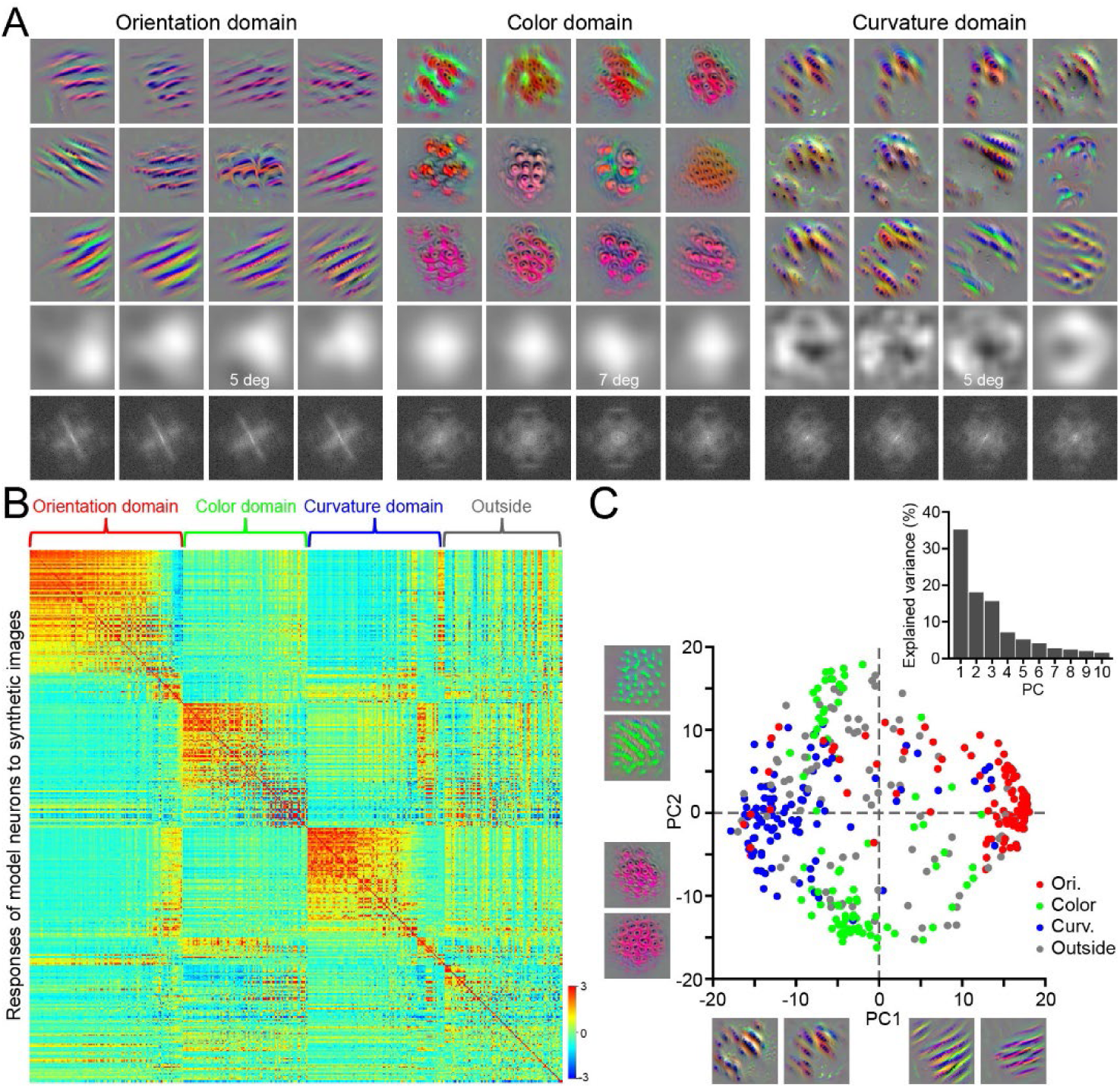
Characteristic SIs in three types of V4 domains. **A**: SIs of V4 cells from three types of functional domains and their spatial features. Left: V4 orientation cells; middle: V4 color cells; right: V4 curvature cells (each group from the same site). Top 3 rows: SIs; row 4: RF of the neuron in row 3, i.e., the “where” layer in the DCNN model. Row 5: spatial frequency map of the feature map in row 3. Images size 5 degrees for orientation and curvature domains, and 7 degrees for color domain. **B**: Response matrix (z-score −3 to 3) of cell models to SI stimuli. X-axis: 385 neurons, divided into 4 categories (Orientation: n=112; Color: n=90; Curvature: n=99; outside (cells not in any domains): n=84). Within each category, neurons are sorted by response strength. Y-axis: SI stimuli, arranged in the same order as the X-axis neurons. **C**: PCA results of the data shown in B, analyzed along the stimulus dimension, showing the distribution of cells in PC1-PC2 stimulus space. PC1 and PC2 are weighted sum of all the SIs, and two example SIs with large weights are shown at each end of each axis. Cells from different functional domains are labeled in different colors; gray cells are from non-domain regions. Inset: variance explained by the first 10 PCs.

We hypothesized that SIs we obtained (e.g. Figure 4A) can be used as effective stimuli in examining V4 neurons, just like gratings were found as effective stimuli in examining V1 neurons. That is, different types of neurons can be maximally differentiated by such SI stimulus sets. To verify this, we used the SIs of 385 V4 cells with EVE > 0.4 as visual input to each cell model, obtaining a 385×385 response matrix. Figure 4B visualizes this response matrix (z-scored) arranged by the cells’ functional domains; the strong responses near the diagonal indicate similar preference among cells of the same type. We performed PCA dimensionality reduction on this 385×385 matrix along the stimulus dimension (Y-axis). The resulting PCs are linear weighted sums of the SIs; the first two dimensions (PC1 and PC2) explained 35% and 18% of the response variance, respectively. We visualized the responses of the 385 cells in PC1-PC2 space (Figure 4C). Cells are colored according to their functional domains. Cells from different functional domains were largely separated, each clustering in different regions of PC space. Importantly, the clusters obtained largely matched the functional domain cell groups, indicating that functional domains identified by ISOI indeed have a cellular basis, rather than a hemodynamic pattern merely reflecting differences in population activity. That is, functional imaging captures the main features of cells. This is confirmed by examining the nature of the PC axes and the SIs with large weights on the PCs (examples shown on axes in Figure 4C). PC1 represents shape (spiky-stubby) information, with orientation and curvature at opposite ends of the axis. PC2 represents color (cool-warm) information, with red and green being separate representations. In the figure, shape-preferring and color-preferring cells are located at the two ends of the X and Y axes, respectively, and are orthogonal to each other, suggesting that these two types of information are represented independently in V4. Similar analysis using the original responses of cells to natural objects also supported the conclusion of independence between shape (spiky-stubby) and color (cool-warm) (Figure S4). Thus, blind-source clustering analysis reveals the similarity of cells within functional domains and the main differences between cells of different functional domains.

### In silico lesioning uncovers distinct input-integration rules for orientation and curvature cells

To understand the neural mechanisms underlying feature selectivity in different V4 neurons, we further studied the DCNN models. The first layer (L1) of the DCNN has 96 convolutional kernels, as visualized in Figure 5A. We classified these kernels into three types, in the order of decreasing orientation features and increasing color features. They are: orientation kernels (1-32), orientation-color mixed kernels (33-67), and color kernels (68-96). For every cell, we “lesioned” each kernel individually to assess its impact on the model’s goodness-of-fit (EVE), and defined the percent loss in EVE as the contribution weight of that kernel to the cell. In Figure 5B, each column shows the weights of 96 kernels (arranged in the order from Figure 5A) for one cell; different columns are different cells (n=385), grouped by their functional domain (orientation, color, curvature). It can be seen that orientation and curvature cells receive their strongest inputs from orientation-color mixed kernels, rather than pure orientation kernels. Color cells, in contrast, receive their strongest inputs from color kernels, while also receiving inputs from the orientation-color mixed kernels (Figure 5B&C). For orientation cells, their preferred orientation was significantly correlated with the orientation of the kernel with the largest weight (Figure 5D), indicating that they inherited orientation selectivity from corresponding L1 orientation kernels. Interestingly, some curvature cells also have orientation preferences (their long-axis orientation), and that orientation is also correlated with the orientation of their main input kernels (Figure 5E).

**Figure 5.**
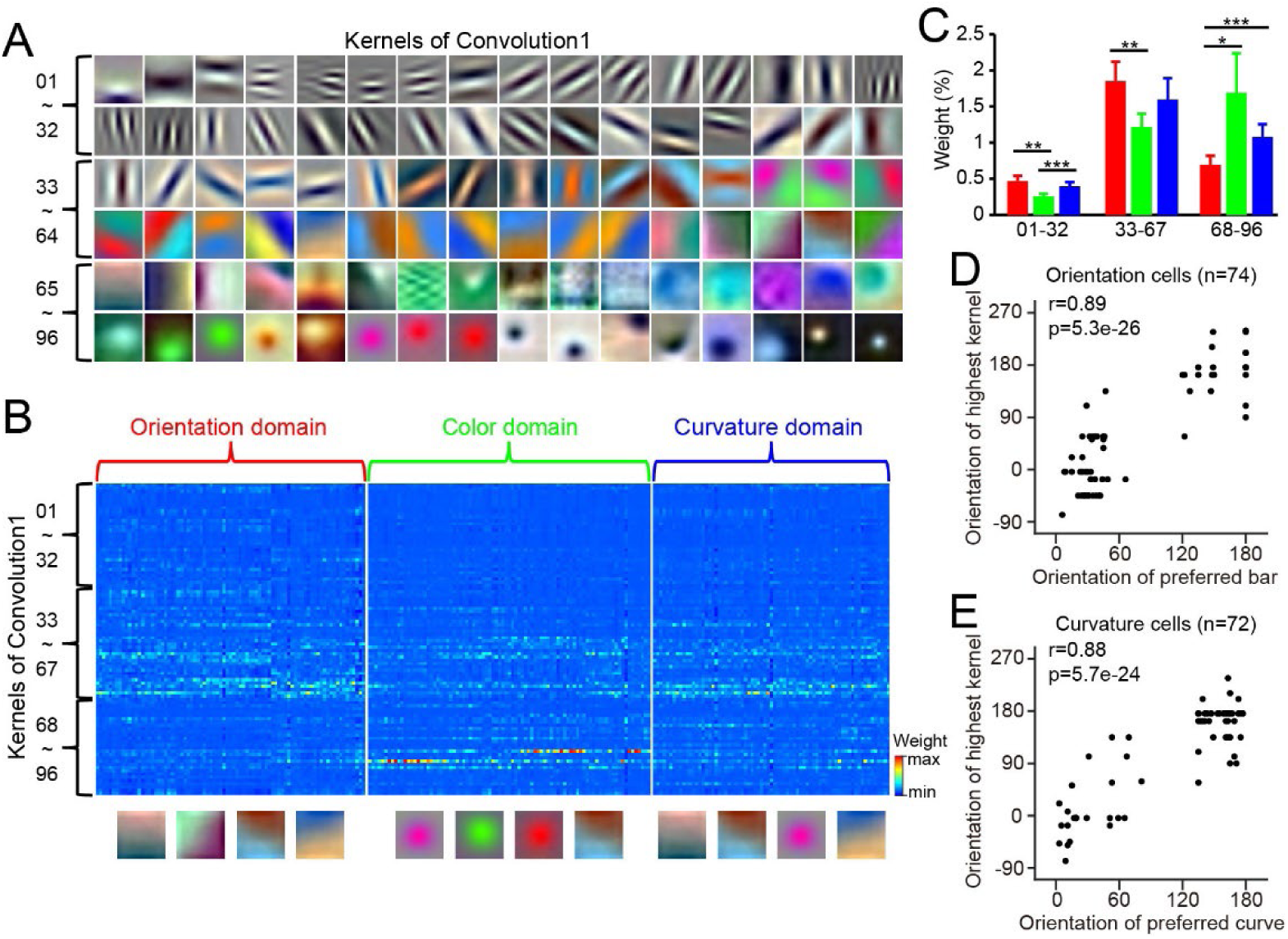
Contribution of DCNN layer 1 kernels to different types of V4 neurons. **A**: Visualization of the 96 convolutional kernels in the first layer of the model. Kernels are sorted in the order of decreasing orientation selectivity and increasing color selectivity, and divided into three categories (1-32: pure orientation; 33-67: orientation-color; 68-96: pure color). **B**: Contribution weights of different kernels (Y-axis) to different cells (X-axis) (percentage impact on EVE accuracy after lesioning a kernel). Kernels are arranged (top to bottom) in the same order as in A. **C**: Average contribution weights of the three kernel categories to each cell category, based on data in B (t-tests, * p<0.05; ** p<0.01; *** p<0.001). **D**: Correlation between the preferred orientation of orientation cells and the orientation of the L1 kernels with the largest weight (r=0.89, Pearson correlation, p=5.3e^-26^). **E**: Correlation between the long-axis orientation of curvature cells and the orientation of the L1 kernels with the largest weight (r=0.88, Pearson correlation, p=5.7e^-24^).

Both orientation and curvature cells process shape information, and the total weight of orientation inputs they receive is comparable (Figure 5C). Yet, they develop very different shape preferences. We analyzed their L1 inputs with different orientation preferences, and how their contribution may make a difference. For each cell we selected their 10 L1 kernels with the highest weight (excluding those without significant orientation, i.e., 68-96 in Figure 5A). For each pair of the kernels, we calculated the difference in preferred orientation (Y-axis of Figure 6A) and the similarity of their impact on the neuron’s response, which is the correlation between the response gradients of the neuron to the 800 stimulus images under perturbation of each kernel (see Methods: Evaluation of Layer 1 kernel efficacy). Analysis revealed that orientation and curvature cells integrate different orientation inputs differently. The negative correlation in Figure 6A indicates that the more different the orientations of two kernel inputs, the more opposite their effects on V4 orientation cells. That is, for V4 orientation cells, inputs with similar orientations have synergistic effects, while inputs with perpendicular orientations have antagonistic effects. In contrast, for curvature cells, the correlation between inputs of different orientations was very weak, close to zero (Figure 6C). Figure 6B&D present comparisons of correlations for similar and perpendicular orientation pairs, showing this difference more clearly. Figure 6E summarizes this finding: although both orientation and curvature cells receive inputs from common orientation kernels, they integrate these inputs differently. For V4 orientation cells, synergy among similar orientations and antagonism among perpendicular orientations enhance their orientation selectivity. While V4 curvature cells lack this antagonism mechanism; they combine inputs of different orientations according to a spatial weighting scheme (weak center, strong periphery) to produce their unique curvature preference.

**Figure 6:**
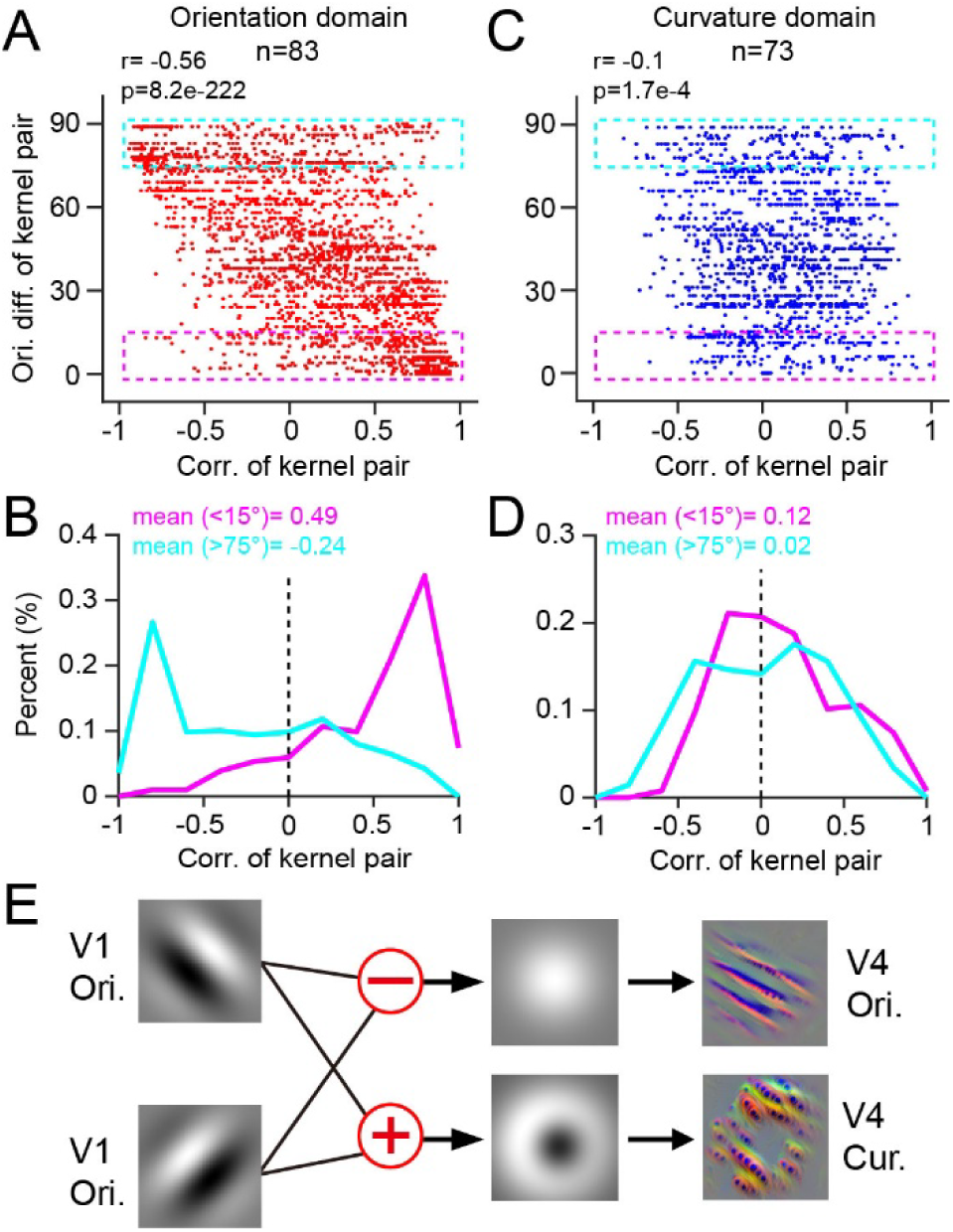
Different integration ways of orientation inputs for orientation and curvature neurons. **A**: For orientation neurons, the orientation difference between L1 orientation kernels (Y-axis) is negatively correlated with the similarity of their contributions to the cell’s response (X-axis). The data in dashed box is quantified in B. **B**: For L1 orientation kernel pairs with similar orientations (differences <15°), their contributions to orientation cells’ responses are positively correlated (pink curve, from pink box in A). For L1 orientation kernel pairs with perpendicular orientations (differences >75°), their contributions to orientation cells’ responses are negatively correlated (cyan curve, from cyan box in A). **C**: Similar to A, but for curvature neurons, showing a weaker correlation (r=-0.1) between orientation difference of L1 orientation kernel pairs and similarity of their contributions to curvature cell responses. **D**: Similar to B, for curvature neurons, showing little differences between kernel pairs with similar orientation preferences and those with perpendicular orientation preferences. **E**: Diagram illustrating the different orientation integrations in V4 orientation and curvature cells.

Similarly, we also analyzed the integration of color inputs in V4 cells. We found that V4 color-preferring cells integrate different color inputs similarly to how V4 orientation cells integrate orientation inputs: similar colors contribute similarly, and opposite colors antagonize each other (Figure S5 A-B). Although curvature and orientation cells also receive color inputs (Figure 5 C-F) and respond significantly more strongly to color SIs than to grayscale SIs (Figure S5 G-H), the antagonism between opposite color inputs is weaker (Figure S5 C-F), and the synergy between similar color inputs is above chance level (Figure S5 C-F). These analyses demonstrate that after building cell models, we can perform a series of experiments on these digital twins that would be difficult to conduct in *in vivo* experiments. Although the results require further validation, model-based work can provide insights for future validation studies.

## Discussion

In this study, we imaged calcium responses of macaque V4 neurons to natural objects, and trained single-cell DCNN models. Modeled neurons formed distinct subgroups in data-driven clustering analysis. Importantly, the main dimensions distinguishing cell groups (PC1 & PC2) coincided with the major mapping features of the known functional maps in V4 (color, orientation, curvature), indicating that ISOI maps can capture the essential characteristics of V4 neurons. Further analysis of the internal connections in different neuron models revealed systematic differences in their inputs, which help to better understand the mechanisms underlying their response characteristics.

### Calcium imaging and modeling reveal complex receptive field features

V4 neurons have complex visual responses and exhibit multi-dimensional selectivity (Tanaka et al. 1986; Desimone & Schein 1987). Traditional parametric measurements of selectivity using visual features have encountered difficulties in fully capturing a cell’s characteristics, or even in determining a cell’s major selectivity dimension. Using calcium signals recorded via two-photon imaging, we successfully built DCNN models of V4 neurons and further obtained SIs for each neuron through these “digital twins”. These SIs typically contain complex, mottled textures and combinations of single or multiple colors (Figure 2D&E, Figure 3B&C, Figure S2A). They resemble neither the artificial visual stimuli commonly used in traditional studies nor natural images. Obtaining such specific images with traditional methods is challenging. The synthesis of SIs in our model was not constrained by natural image priors; the sole goal was to maximize the neuron’s response. The algorithm freely combines features that are effective in the neural representation space. The grayscale versions of these results (Figure S5G) resemble those of recent studies from other groups (Bashivan et al. 2019; Willeke et al. 2026), confirming the validity of our work. The novelty of our work lies in using color images as visual stimuli, training models that simulate cells’ color responses. We not only found differences in V4 cells’ color preferences (e.g., Figure 3), but also discovered orthogonal coding between color selectivity and shape selectivity (Figure 4C) and their spatial segregation on the cortex (Figure 4C, Figure S3). This suggests that although these two features coexist in single cells, they remain relatively independent (weak interaction), and do not exhibit overt feature binding. This property continues the characteristics of V1 (Garg et al., 2019; Wang et al., 2026) and V2 (Friedman et al., 2003, our own unpublished observation) cells.

We found that most V4 cells have one major feature, which can be identified from their SIs. The SIs of orientation, color, and curvature cells are clearly distinct. Groupwise, neuron’s major feature becomes even more evident (Figure 3, Figure S3). Although previous studies have also described V4 neurons using features such as color, orientation, and shape (Schein et al. 1982; Tanaka et al. 1986; Desimone & Schein 1987), such characterization was often inaccurate due to the limited number of stimuli. Our results show that optical imaging maps indeed capture the major features of V4 cells. Although this correspondence is not always necessary (e.g., color-selective neurons could share the same orientation preference and thus form an orientation domain), our results did not show this. Our results in V2 are similar (unpublished observation).

Our knowledge of V4 neuronal responses largely came from electrophysiological recordings, and recent DCNN modeling has also been based on electrophysiological single-unit responses (Bashivan et al. 2019; Willeke et al. 2026). Although two-photon calcium imaging has limited temporal resolution, our results show that calcium signals, as electrical signals, can also be used for DCNN model training and achieve relatively good model fit. Models trained with calcium signals successfully captured neuronal differences in receptive field size, input complexity, and complex spatial structure of SIs (color and texture). In addition, calcium imaging has the advantage of preserving the spatial information of cells, providing direct cellular-level evidence for the functional architecture of V4 (Figure 4). The consistency between these functional structures and ISOI functional maps further supports the validity of the models.

The digital twins of V4 neurons allowed us to try experiments that difficult to perform in animals. By lesioning first-layer convolutional kernels, we obtained their contribution weights to the neurons under study (Figure 5B). We found that both orientation and curvature neurons inherited orientation selectivity from their inputs (Figure 5 D&E). For V4 orientation cells, this was expected (Fang et al. 2022; Cai et al. 2023). For curvature cells, however, this was not a foregone conclusion, as V4 could in principle generate curvature receptive fields de novo solely through the spatial arrangement of inputs. Moreover, V4 curvature cells and orientation cells are opposed along the shape dimension (Figure 4C). We found that curvature neurons integrate inputs of different orientations in a manner distinct from orientation neurons (Figure 6E). Thus, orientation and curvature cells share common inputs but adopt different integration strategies, producing dramatically different cellular response characteristics. Given the similarity between the DCNN and the hierarchical structure of the ventral pathway, these findings from virtual cells are plausible.

### Direct evidence for functional architecture in V4

Our work reveals the functional architecture in V4 at the cellular level through both cortical spatial distribution of SIs and PCA clustering. First, the cortical spatial distribution of neuronal SIs showed clear clustering features (Figure 3, Figure S3). Adjacent cells had similar SIs in shape, color, and texture aspects. Within the spatial extent of a single two-photon imaging site (∼0.8mm), the SIs were similar, with gradual shifting preferences, eg. hue (Figure 3A) or preferred angle (Figure S3), forming smaller sub-domains. SIs from different sites differed substantially, and their types could be easily distinguished from the SI features. SI similarity indicates that cells have similar preferences in high-dimensional stimulus space (Willeke et al. 2026; Wang et al. 2024; Burg et al. 2024). This similarity is weaker in the low-dimensional space tested by conventional stimuli, leading to the conclusion that similarity among neighboring cells is low (e.g., Namima et al. 2025). At the population activity level, subtle differences in low-dimensional space can be amplified in contrast maps, such as functional maps obtained with ISOI (e.g., Ghose & Ts’o 1997; Tanigawa et al. 2010). These maps typically reveal a single feature dimension, reflecting the projection of high-dimensional preferences onto a particular dimension. Additionally, recent studies have found clustering of V4 neurons through model computations (Hatanaka et al. 2022) or revealed cortical functional segregation through population calcium imaging (Wang et al. 2024).

Another line of evidence for V4 functional architecture comes from purely data-driven clustering analysis (PCA), which revealed distinct cell clusters that are spatially segregated on the cortex. The main differences distinguishing different V4 neurons came from their responses to shape (PC1) and color (PC2). As shown in Figure 4C, on the PC1 shape dimension, curvature and orientation are opposed and represented by two independent cell populations, namely orientation cells and curvature cells, which have no systematic differences in their PC2 coordinates. On the PC2 color dimension, cells preferring cool colors and warm colors separate along this dimension, with no difference in PC1. These phenomena reflect the independence between these two feature types. Importantly, these two major visual dimensions correspond exactly to the most prominent functional map types previously revealed in V4 population activity, including color maps (Conway et al. 2007; Tanigawa et al. 2010), orientation maps (Ghose & Ts’o 1997; Tanigawa et al. 2010), and curvature maps (Tang et al. 2020; Hu et al. 2020; Jiang et al. 2021). Thus, our work not only provides direct cellular-level evidence for functional maps, but also establishes a correspondence between functional map features and the major features of single cells. It demonstrates that the functional architecture obtained with blood-oxygen signal imaging indeed reflects the major characteristics of the V4 neuronal population.

The functional architecture of V4 is closely related to its inputs. Previous studies have found that different V4 regions receive parallel projections from V2 (Zeki & Shipp 1989; Nakamura et al. 1993; DeYoe et al. 1994; Felleman et al. 1997; Xiao et al. 1999). Our previous work showed that V4 color domains predominantly receive input from V2 thin stripes, while orientation domains predominantly receive input from pale stripes (Fang et al. 2022). Further upstream, V2 color and shape (orientation) modules also receive parallel projections from V1 blobs (color-dominant) and interblobs (orientation-dominant) (Livingstone & Hubel 1983; Sincich & Horton 2002). Therefore, our confirmation of the existence of functional architecture in V4 supports the view that the ventral pathway processes color and shape information in parallel. The functional architecture of V4 is one link in the information processing chain of the entire visual ventral pathway, reflecting the cortex’s strategy of parallel processing of visual information.

### Limitations and novelty of this work

This work is based on calcium signals. Compared with electrical signals, neurons’ calcium responses are slower and have lower signal-to-noise ratio, so each trial yields less cell-based information, requiring longer recording times. We found that for more complex V4 neurons, the goodness-of-fit was significantly lower than for V1 (Figure 2C, Figure S2B-E), indicating that more training samples are needed to measure V4 neurons. Second, we did not measure in the direction perpendicular to the cortical depth; imaging depth was limited to 155–295 μm, corresponding approximately to cortical layers 2-3. We have no information about deeper neurons, and we cannot directly support the existence of columnar structures. Third, in this study, we selected orientation, curvature, and color functional domains with strong functional maps. In addition to these three types, other studies have reported other types of functional architecture, including size (Ghose et al. 1997; 2017), motion direction (Li et al. 2013; Fang et al. 2022), SF (Lu et al. 2018; Zhang et al. 2023), 3D shape (Srinath et al. 2021), and border ownership (Franken et al. 2021). For these functional domains not covered here, the approach of two-photon imaging combined with DCNN modeling used in this study should also be applicable.

This work has innovations both methodologically and scientifically. Methodologically, this is the first study to combine two-photon calcium imaging and DCNN modeling to study macaque visual neurons, validating the feasibility and advantages of this approach. We also used color images, characterizing the color processing properties of V4 cells and discovering the orthogonal relationship between color and shape. Another methodological feature of our work is the combination with ISOI, establishing a precise correspondence between population activity functional domains and single-cell functional clustering, providing direct evidence for their consistency. In addition, we leveraged the unique advantages of digital twins to investigate the sources of neuronal inputs, obtained preliminary results that are meaningful for future work. These methodological innovations yielded new results. We found that functional architecture has a cellular basis and is built upon the major response features of cells, indicating that cortical functional architecture has important functional significance.

Different visual features are represented in a segregated manner both at the single-cell level and at the population functional architecture level, suggesting that visual features are processed primarily in a segregated fashion in V4. These findings have important implications for our understanding of V4 function.

## Acknowledgement

This work was supported by the National Natural Science Foundation of China (32271079, 31625012, 31530029, and 31800870) and the Brain Science and Brain-like Intelligence Technology –National Science and Technology Major Project (2022ZD0204600). We thank lab members for providing technical assistance.

## Author contributions

R.T., H.H., and H.L. conceptualized the project. R.T., W.Z., R.Z. and J.W. performed the experiments. R.T., Q.Z., C.D. and H.L. analyzed the data. R.T., Q.Z., G.W., C.D., H.H. and H.L. wrote the manuscript

## Materials and Methods

### Animals

All animal procedures in this study were performed in accordance with the National Institutes of Health Guidelines and were approved by the Institutional Animal Care and Use Committee of the Beijing Normal University with protocol number: IACUC(BNU)-NKCNL2016-06. Three hemispheres from three animals (two rhesus macaques, one cynomolgus macaque, all male) were studied. All hemispheres underwent both intrinsic signal optical imaging (ISOI) and two-photon calcium imaging (2P imaging). These animals were also used for other studies (Zhang et al. 2025). Basic experimental procedures were similar to those previously published by our laboratory (Tang et al. 2020).

### Surgical Procedures

Animals were initially anesthetized with ketamine (10 mg/kg) in their home cages. They were then transferred to the laboratory, where the head was fixed in a stereotaxic apparatus. Under isoflurane (1.5%-2.5%) anesthesia with a ventilator, a craniotomy (22 mm diameter) was performed. The center of the craniotomy was 18-35 mm from the midline and 15-25 mm from the occipital ridge, targeting V1 or V4. The dura mater was opened, and visual areas V1, V2, and V4 were identified based on the locations of the following sulci: lunate, superior temporal, inferior occipital. Intrinsic signal optical imaging was first performed on the exposed visual cortex. Based on the functional maps obtained, targeted virus injections were then made into the identified functional domains (10-15 injection sites per chamber, with approximately 3/4 of injection sites located at the centers of functional domains and the remainder in non-domain areas). The viral vector was AAV9.Syn.GCaMP6S.WPRE.SV40 (CS1282, titer 3.34e13 GC/ml, Addgene). Injection depth was 500 μm, and injection volume was 500 nL per site. After virus injection, the cranial window was sealed, and a second surgery was performed 6 weeks later to implant a sealed optical window (13 mm diameter, Li et al. 2017). All subsequent ISOI and 2P were performed through this sealed optical window, with imaging sessions spaced 7-10 days apart.

### Intrinsic Signal Optical Imaging

Intrinsic signal optical imaging on the day of surgery was performed without an imaging window; the exposed cortex was covered with a coverslip and flattened, with agar filling the space. After the imaging window was implanted, imaging could be performed directly through the window. During imaging, anesthesia was switched from isoflurane to intravenous anesthetics. Two anesthesia protocols were used in this study: Protocol 1: induction with propofol 2.0 mg/kg + sufentanil 0.15 μg/kg, maintenance with propofol 2.0 mg/kg/h + sufentanil 0.15 μg/kg/h (case 1); Protocol 2: induction with ketamine 5.0 mg/kg, maintenance with ketamine 5.0 mg/kg/h (cases 2 and 3). Vecuronium bromide (induction 0.25 mg/kg, maintenance 0.05 mg/kg/h) was also infused intravenously to maintain eye stability. Animal heart rate, body temperature, blood oxygen, and end-tidal CO₂ levels were continuously monitored to assess physiological status and anesthesia depth. Anesthetic infusion rates were adjusted accordingly during imaging. The pupils were dilated with atropine (0.5 mg/ml), and the animal’s eyes were focused on a screen 57 cm in front using contact lenses of appropriate curvature. Cortical reflectance changes (intrinsic blood-oxygen signals) were recorded at 4 Hz using an Imager 3001 system (Optical Imaging, Inc., Germantown, NY, USA). Images were 540 × 654 pixels, corresponding to a cortical area of 15.5 × 19 mm. In each trial, visual stimuli were presented for 3.5 s. Imaging started 0.5 s before stimulus onset and ended at stimulus offset, for a total of 16 frames in 4 s. Inter-stimulus interval was approximately 6 s.

### Two-Photon Calcium Imaging

Animal preparation and anesthesia were the same as for ISOI. The microscope used for 2P was a Prairie Ultima IV (In Vivo) 2P microscope (Bruker Nano), with a Chameleon laser (Coherent). Laser wavelength for was set to 980 nm, with power of 30-45 mW. The objective was a 16x water immersion lens (0.8-N.A., Nikon), corresponding to an imaging area of 830 × 830 μm (except for site 4, which was 415 × 415 μm). The scanning mode was Galvo mode, with a scan rate of 1.3 Hz and image size of 512 × 512 pixels. The onset time of each visual stimulus was always aligned with the 2P scanning phase, i.e., the next visual stimulus was presented when the scan began for the next frame.

Because the microscope objective was vertical, during 2P, we used a custom-made rotatable mount to tilt the animal’s head approximately 70 degrees so that the chamber glass was parallel to the objective lens, and the animal was lying on its side. The stimulus screen was also rotated by the same angle. In this study, 2P was performed at 8 sites across 3 hemispheres (Figure S1). Imaging depths were ranging from 155–295 μm from the cortical surface. Only one depth was imaged per site.

### Visual Stimuli for Intrinsic Signal Optical Imaging

Visual stimuli were generated by a ViSaGe system (Cambridge Research Systems Ltd.) and presented on a 21-inch CRT monitor (SONY CPD-G520) placed 57 cm in front of the animal. Both eyes of the animal were open. The monitor luminance was gamma corrected and running with a refresh rate of 100 Hz. Full-screen sinusoidal gratings were used to obtain orientation maps and color maps. Grating stimuli included 2 spatial frequencies (0.25 and 1 cycle/deg) and 8 motion directions (0-315°, 45° intervals), drifting at a temporal frequency of 4 cycles/s. The mean luminance of both black-white gratings and red-green gratings was 28.92 cd/m². Full-screen shape stimuli (circle & triangle) were used to obtain curvature maps (Tang et al. 2020). Each shape stimulus was 2.5 degrees and was arranged in a grid with 3-degree spacing. The luminance of the bright lines was 111.2 cd/m², line width of 0.2 degrees; the background luminance was 20.57 cd/m². In each trial, the full-screen stimulus drifted in one of 8 random directions (0-315°, 45° intervals) at a speed of 4 deg/s.

### Visual Stimuli for Two-Photon Calcium Imaging

Visual stimuli were generated by a ViSaGe system and presented on a 21-inch LCD monitor (DELL E1913Sf) placed 57 cm in front of the animal. The monitor refresh rate was 60 Hz, and was gamma corrected. Stimulus patches had maximum luminance of 80.82 cd/m² and were presented on a gray background with luminance of 40.41 cd/m². All stimuli were presented to the animal’s left eye, with the right eye closed.

To determine the receptive field (RF) location of neurons, we first manually moved a circular patch of drifting square-wave grating (1-3 degrees) across the screen while observing cortical fluorescence responses to estimate the approximate RF location. Then, centered on that estimated location, we presented visual stimuli at random locations within a 5×5 grid. Stimuli were three shapes (square-wave grating, triangle, circle) sized 1-3 degrees, drifting at one of 4 directions (45-315°, 90° intervals) with a speed of 4 deg/s. Each stimulus was presented for 1.5 s, with an inter-stimulus interval of 0.8 s. After stimulus presentation, population RF location and size were estimated online (Gaussian fit, threshold set at 0.5× Gaussian height).

To study neuronal responses to natural visual stimuli, we constructed a natural object stimulus set consisting of 800 color images without backgrounds, selected from 450 object categories (www.freepngs.com). For 5 V4 sites, the stimulus set also included 50 artificial shape stimuli, including gratings, short bars, and simple geometric shapes. All stimuli were presented without luminance adjustment against a gray screen. Stimuli size was set to 5 degrees for V1 imaging and 80% of the RF size for V4 sites. Stimuli were presented in a random order, each for 2 seconds, with an inter-stimulus interval of 3-4 seconds. To avoid adaptation, during the first second of stimulus presentation, the stimulus size gradually increased from 80% to 95% of RF size, and during the second second, it gradually decreased back to 80% of RF size. After all stimuli had been presented once, RF location was remeasured before the next trial. Each cortical site was imaged in at least 3 experiments. We collected 5 trials of responses to natural object in each of the first two experiments. After that the data were used for DCNN model training. The trained DCNN model was used to obtain synthetic images (SI) for each neuron (see Neuronal DCNN Encoding Model). These SIs were tested in the third experiment, along with the regular stimulus set.

### ISOI Data Analysis – SVM Maps

ISOI data were primarily used to calculate cortical functional maps (e.g., Figure 1B-D). Similar to previous studies (Tang et al. 2020), functional maps were obtained using support vector machine (SVM) methods. ISOI data analysis was performed using MATLAB 2017 (The MathWorks, Natick, MA). We first calculated a response map (ΔR/R) for each stimulus in each trial. ΔR/R = (R_8-16_ – R_1-4_)/R_1-4_, where R_8-16_ is the average of frames 8–16, and R_1-4_ is the average of frames 1–4. Based on these response maps, we then computed SVM maps by applying an SVM algorithm to classify response maps of two stimulus types, obtaining the weight of each pixel in the overall classification. The weight value of a pixel indicated how much that pixel contribute to distinguish the two stimulus types, with the sign indicating preference for one stimulus type over the other. The Matlab SVM program was provided by Chih-Jen Lin (LIBLINEAR: A Library for Large Linear Classification, 2008; available at https://www.csie.ntu.edu.tw/~cjlin/liblinear/).

For each SVM map, Gaussian low-pass filtering (size=5, std=1) was first applied. The image was then segmented into cortical area and background (including blood vessels and areas outside the chamber) based on a vascular map (green channel). The cortical area was normalized (z-scored) using the standard deviation of gray values in the background area, and ±2 standard deviations were used as the threshold for determining significant domains. Additionally, domain size was required to be >50 pixels. Two-photon images from each site were also aligned with the vascular map, enabling the assignment of domain type to each cell within the OI maps.

### 2P#Data Analysis

Two-photon data analysis was also performed using MATLAB 2017. Image alignment and cell extraction algorithms were the same as in our previous study (Tang et al. 2020). Images from each session at the same site (including data from different days) were aligned to a common template (the average of the first 1000 frames of the first session) to correct for displacement during imaging (typically <20 pixels). Cell ROIs for a given site were extracted based on imaging data from the first day (typically 5 trials). For each session, the image sequence was first segmented by the onset time of each stimulus. Image sequences corresponding to all stimuli were then averaged across the stimulus dimension. The average of the two frames immediately before stimulus onset was taken as the baseline image, and the average of the 2nd and 3rd frames after stimulus onset was taken as the response image. The difference image (response minus baseline) was used to identify responsive cells. A cell was identified if a contiguous region of >30 pixels in a difference image had gray values greater than the mean plus 1.5 standard deviation of the entire image. Cell ROIs obtained from all sessions on that day were then merged, and overlapping ROIs were removed, yielding the final set of cell ROIs for that site. Subsequent data from that site used the first day’s cell ROI set. Once cell ROI were determined, we calculated each cell’s response amplitude to each stimulus as the fluorescence change ratio (ΔF/F_0_), where ΔF = F–F_0_, F is the mean gray value of the cell in the response image, and F_0_ is the mean gray value of the cell in the baseline image. If a cell’s ΔF/F_0_ response amplitude (averaged across trials) for all stimuli in the experiment did not exceed 0.3, that cell was excluded from subsequent analysis. The trial-averaged responses of neurons to each visual stimulus were used for subsequent PCA analysis and DCNN model training and test.

### PCA

Responses of each neuron to different stimuli were first z-scored. The response matrix of the neuronal population was then centered (mean subtracted) along the analysis dimension. Singular value decomposition (SVD) was then performed on the response matrix to obtain PCs and the coordinates of samples on each PC. For PCA on SI responses, the response matrix was replaced by neuron models’ responses to SI stimuli and the same procedures were performed.

### DCNN Model

The DCNN model used in this study was adapted from the model in Bashivan et al. 2019. The model includes pretrained convolutional layers of AlexNet, a “where” layer (corresponding to the RF), and a “what” layer (Klindt et al. 2017) (Fig. 2A). It establishes a mapping from the stimulus image to the pretrained AlexNet feature space and then to the neuronal response space. Unlike the original study (Bashivan et al. 2019), stimulus images were not subjected to fisheye transformation before used in the model. Model fitting in this study consisted of two stages. The first stage was consistent with the literature: the kernels of the pretrained AlexNet convolutional layers were kept fixed, and only the “where” and “what” layers were trained until model convergence. The second stage, which we added, involved fine-tuning the kernels of the pretrained AlexNet convolutional layers along with the “where” and “what” layers until model convergence. This added fine-tuning stage significantly improved the fit accuracy (EVE) of the encoding model (by 0.05–0.1). The loss function for model training consisted of four parts: the mean squared error between actual and predicted neuronal responses: ℒ*_MSE_,* a Laplacian regularization term ℒ_lap_ for the spatial filter, to reduce unnecessary edges in the spatial filter, a regularization term ℒ_lap_ for the spatial filter L2, and a regularization term ℒ*_c_* for the channel filter L2.

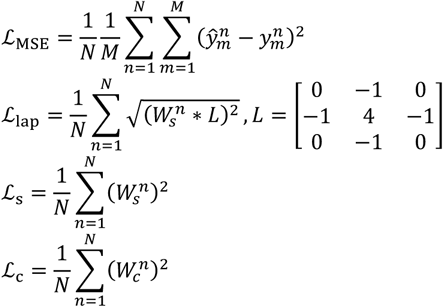

The overall loss function is the sum of these four:

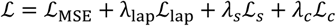

In above functions, N is the total number of neurons, and M is the total number of stimulus images. λ_lap_, *λ_s_*, *λ_c_* are the weight coefficients for parts 2, 3, and 4, respectively. These hyperparameters in the model loss function needed to be optimized during training (see Model Training and Evaluation).

### Selection of Pretrained AlexNet Convolutional Layers

Due to different levels of complexity for V1 and V4 neurons, we used different numbers of AlexNet convolutional layers in modeling V1 and V4 neurons. To quickly select the optimal convolutional layers for each site, we first used sparse random projections (Li et al. 2006) to reduce the dimensionality of the features from each AlexNet convolutional layer. The number of dimensions after reduction was determined by the Johnson-Lindenstrauss lemma based on the number of samples. Then, using a ridge regression model, we used the dimension-reduced feature sets to predict neuronal responses, and used two-fold cross-validation to select the optimal regularization coefficient alpha for each neuron. Finally, based on the average fit accuracy across all neurons in that site, the layer with the highest accuracy was selected as the feature output layer of AlexNet in the encoding model. The optimal AlexNet convolutional layers for the first three V1 sites were layer 1, 2, and 2, respectively. Site 3 was located at the V1/V2 border and may have included a small number of V2 cells. The optimal layer for all five V4 sites was layer 3. These selection results reflect the hierarchical correspondence between AlexNet and the primate visual ventral pathway, consistent with previous findings (Güçlü et al. 2015).

### Single-Site and Multi-Site Models

In the single-site model, only neurons from the same site were trained together. In this case, the convolutional layer parameters of the encoding model were shared across these neurons, while the “where” and “what” parameters were optimized independently for each neuron. The model was initialized for neurons from a new site. The results in Figures 2–4 are based on such a single-site model. To further compare the differences in information processing mechanisms among V4 neurons from different functional domains (mostly distributed across different sites), we additionally constructed a multi-site model. During multi-site model training, neurons from different sites were trained together. The convolutional layer parameters were shared across all neurons, while the “where” and “what” parameters were still optimized independently for each neuron. The results in Figures 5–6 are based on the multi-site model. The overall prediction accuracy of the multi-site model was slightly lower than that of the single-site model.

### Model Training and Evaluation

From the 800 natural stimuli and their corresponding neural responses, 720 were randomly selected to form the training set; the remaining 80 natural stimuli and 50 artificial stimuli (for V4) and their corresponding neural responses formed the test set. During model training, a grid search was performed over the hyperparameters in the loss function: λ_lap_ ∈ [0.01,0.1,1], *λ_s_* ∈ [0.01,0.1,1], *λ_c_* ∈ [0.01,0.1,1] on the test set to select the hyperparameter combination with the highest fit accuracy. The learning rate was set to 1e-4, batch size was set to 50, and the Adam optimizer was used for optimization. An early stopping strategy was used to prevent overfitting: training was stopped if the MSE did not decrease by more than 5e-5 for 500 consecutive iterations.

In this study, the explained variance explained (EVE) was used as a quantitative measure of model fit accuracy.

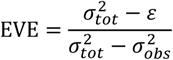

where *ε* is the mean squared error between actual and predicted values, *σt_ot_*^2^ is the variance of neuronal responses across different stimuli, and *σ*^2^*_obs_* is the mean across stimuli of the variance of responses to the same stimulus.

### Synthetic Image (SI)

To visualize the features encoded by each neuron (Bashivan et al. 2019), we used the trained model to generate SIs for each neuron. A gradient ascent approach was used to optimize stimulus images in the pixel space. The loss function was as follows:

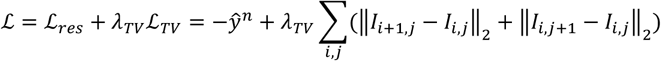

The first term aims to maximize the predicted response of the target neuron, and the second term is a total variation regularization term to prevent excessive high-frequency noise in the image, with *λ_TV_* = 500. Image pixel space was initialized with random noise drawn from a uniform distribution. The learning rate was set to 0.01, and 200 iterations were performed. We also generated a grayscale version of SI by constraining values in RGB channels to be equal.

### Evaluation of Layer 1 Kernel Weights

We lesioned a kernel to evaluate its contribution to a modeled neuron. In the trained multi-site model, each kernel in the first convolutional layer was set to zero one at a time, and the EVE of the encoding model on the test set was recalculated. The ratio of the EVE loss caused by lesioning a given kernel to the total EVE was used as the weight index of that kernel for the modeled neuron.

### Evaluation of Layer 1 Kernel Efficacy

We investigated the influence of each Layer 1 kernel on the responses of V4 model neurons to stimuli. In the trained multi-site model, for the m^th^ stimulus image, the feature map *x* ∈ *R^H^*^×*W*×*C*^ after passing through the Layer 1 convolutional layer was computed, where *H*, *W*, *C* are the height, width, and number of channels of the feature map, respectively. The model outputs the predicted response 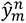 of the n^th^ neuron to this stimulus. The gradient matrix of this neuronal response with respect to the feature map was computed as 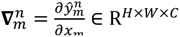, then averaged across spatial dimensions to obtain the response gradient vector of this neuron for the m^th^ stimulus with respect to Layer 1 across C channels: 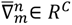. This vector represents the influence magnitude (efficacy) of each of the C channels on the neuron’s response to the m^th^ stimulus. Repeating this process for all M stimulus images yielded the response gradient matrix for the n^th^ neuron across M stimuli with respect to the C channels of Layer 1: 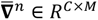. The row vector for the c^th^ channel (kernel) represents the efficacy vector of that kernel for the current stimulus set on this neuron.

To analyze the difference in influence of any pair of Layer 1 kernels on the response of the n^th^ neuron, we calculated the Pearson correlation coefficient between the efficacy vectors of that pair of kernels. A positive correlation coefficient indicates that the pair of kernels contributes to the neuronal response in the same direction (synergistic effect); a negative correlation coefficient indicates that they contribute in opposite directions (antagonistic effect).

### Definition of Layer 1 Kernel Orientation

To determine the orientation preferences of the 96 Layer 1 kernels, they were first manually sorted in order of decreasing orientation features and increasing color features (Figure 5A). These kernels were classified into three categories: kernels 1-32 as orientation kernels, kernels 33-67 as orientation-color mixed kernels, and kernels 68-96 as color kernels. The raw size of Layer 1 kernels is 11×11×3, where the third dimension is the color dimension. For kernels 1-67, each kernel was first normalized to 0-255, then resized to 101×101×3 (bicubic interpolation), and then converted to grayscale. A Gabor equation was then fitted to this grayscale image to obtain the preferred orientation of the kernel.

### Definition of Layer 1 Kernel Hue

To determine the hue preferences of the 96 Layer 1 kernels, each kernel was first converted from RGB to HSV format, and the distribution of hue (0-360) across the 121 pixels was examined. Of these, 24 kernels showed a unimodal distribution; these 24 kernels belonged to kernels 68-96 and were defined as pure color kernels. A Gaussian function was fitted to the hue distribution curve to determine the center of the hue distribution, which was taken as the preferred hue of the kernel.

## Supplementary Figures

**Figure S1.**
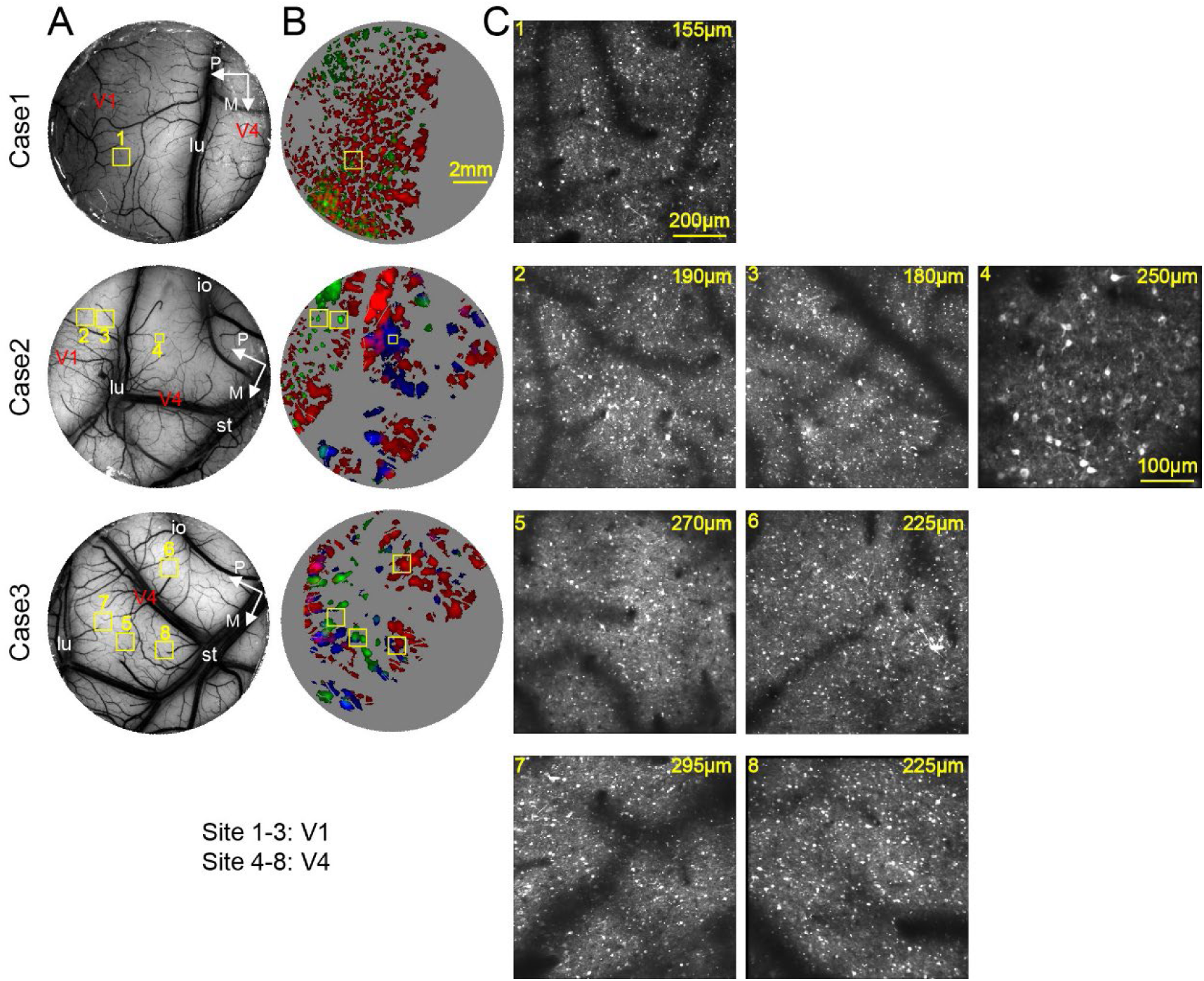
All chambers (n=3) and 2-photon imaging sites (n=8) investigated in this study. (for Figure 1) **A.** Chamber images showing ISOI imaging region and large blood vessel over major sulci: lu, lunate sulcus; st, superior temporal sulcus; and io, inferior occipital sulcus. Yellow square frames indicate regions studied with 2-photon calcium imaging. **B.** Layout of three types of functional domains obtained from ISOI. Red: Orientation domain; Green: Color domain Blue: Curvature domain. Scale bar in case 1 applies to all panels in A and B. **C.** Calcium images from the yellow-framed regions in A and B, obtained with 2-photon calcium imaging. Site numbers are labeled on the top-left corners. Site 8 is also presented in Figure 1F. Each site was imaged from one depth (labeled on top right corner). Scale bar in site 1 applies to all imaging sites except for site 4, which used a zoom.

**Figure S2.**
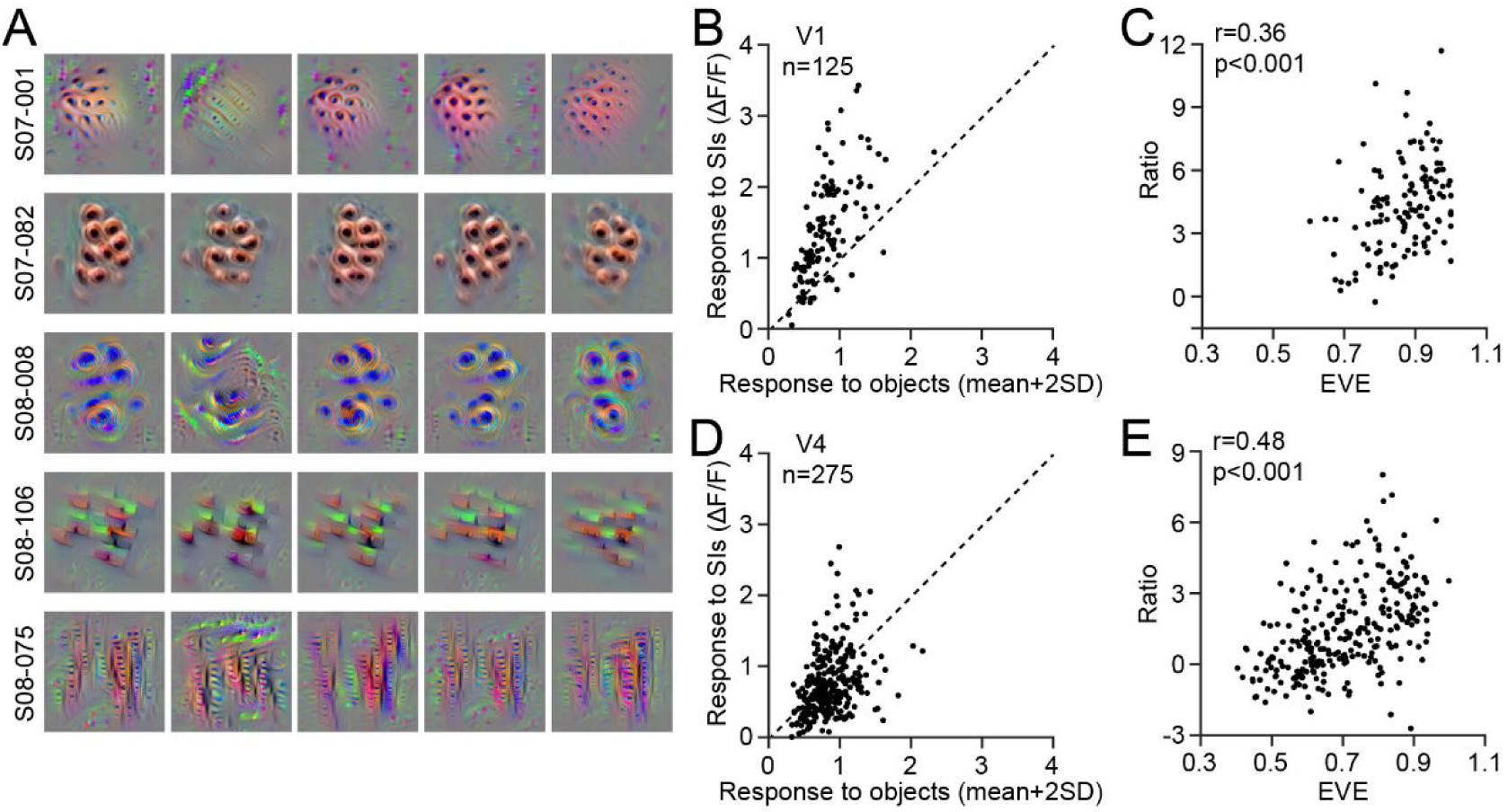
Response of neurons to SIs. (for Figure 2) **A.** SIs of 5 example V4 neurons (5 rows). Each neuron was trained 5 times with partially different training dataset (5-fold training) and different initialization (5 columns). **B.** Response of V1 neurons to their SIs (Y axis) and to natural objects (X axis, mean+2SD) are positively correlated. **C.** Models’ goodness-of-fit is correlated to cells’ response to SI (Y axis, calculated as Ratio = (resp to SI-mean of resp to objects)/std of resp to objects). **D.** Similar to B, but for V4 cells. **E.** Similar to C, but for V4 cells.

**Figure S3.**
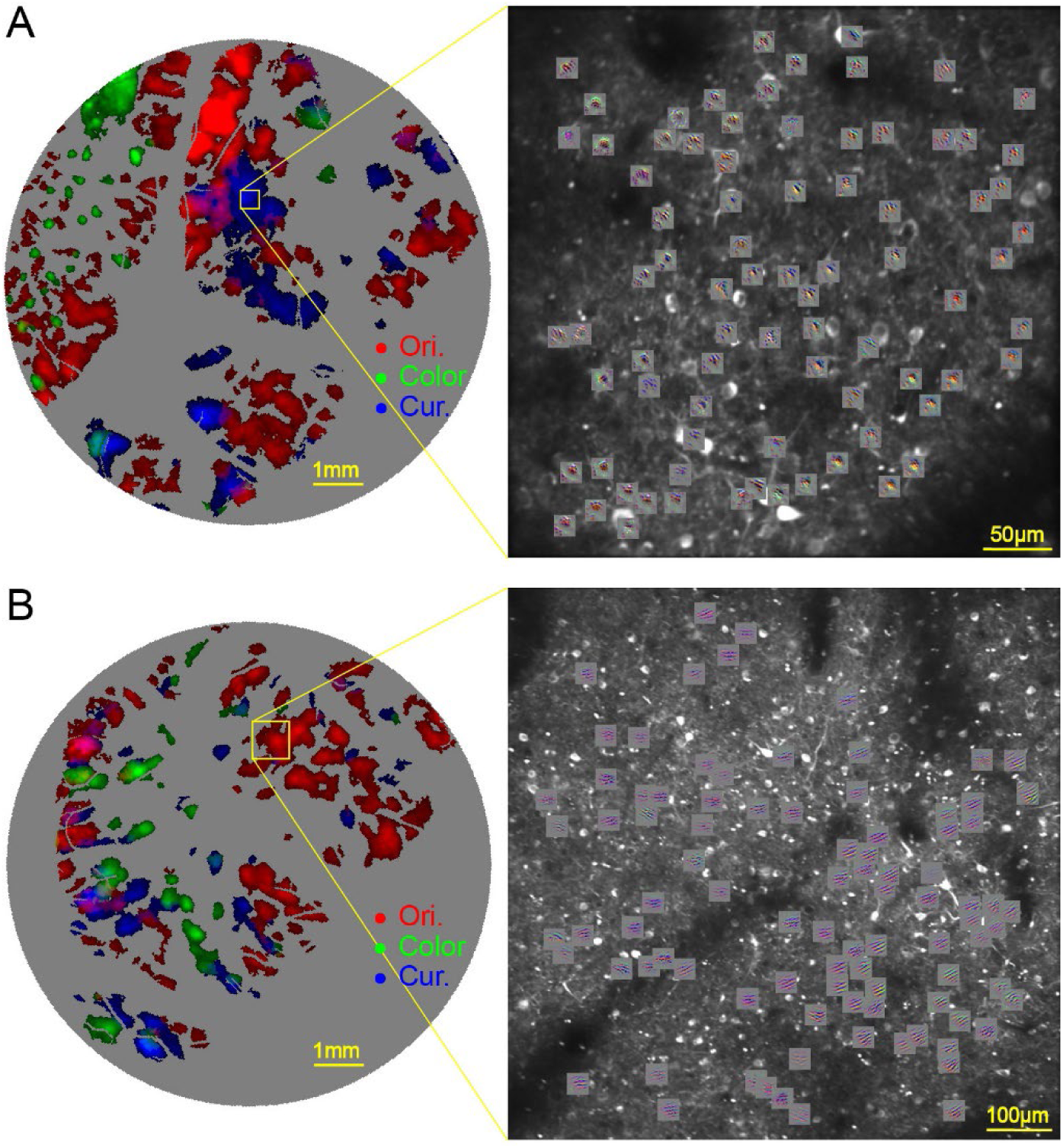
Maps of cellular SI in two example sites (for Figure 3) **A.** Site 4 in Case 2 is located in a curvature domain. Neurons’ SIs in this region exhibit similar features. **B.** Similar to A, but for site 6 in Case 3, which is located in an orientation domain.

**Figure S4.**
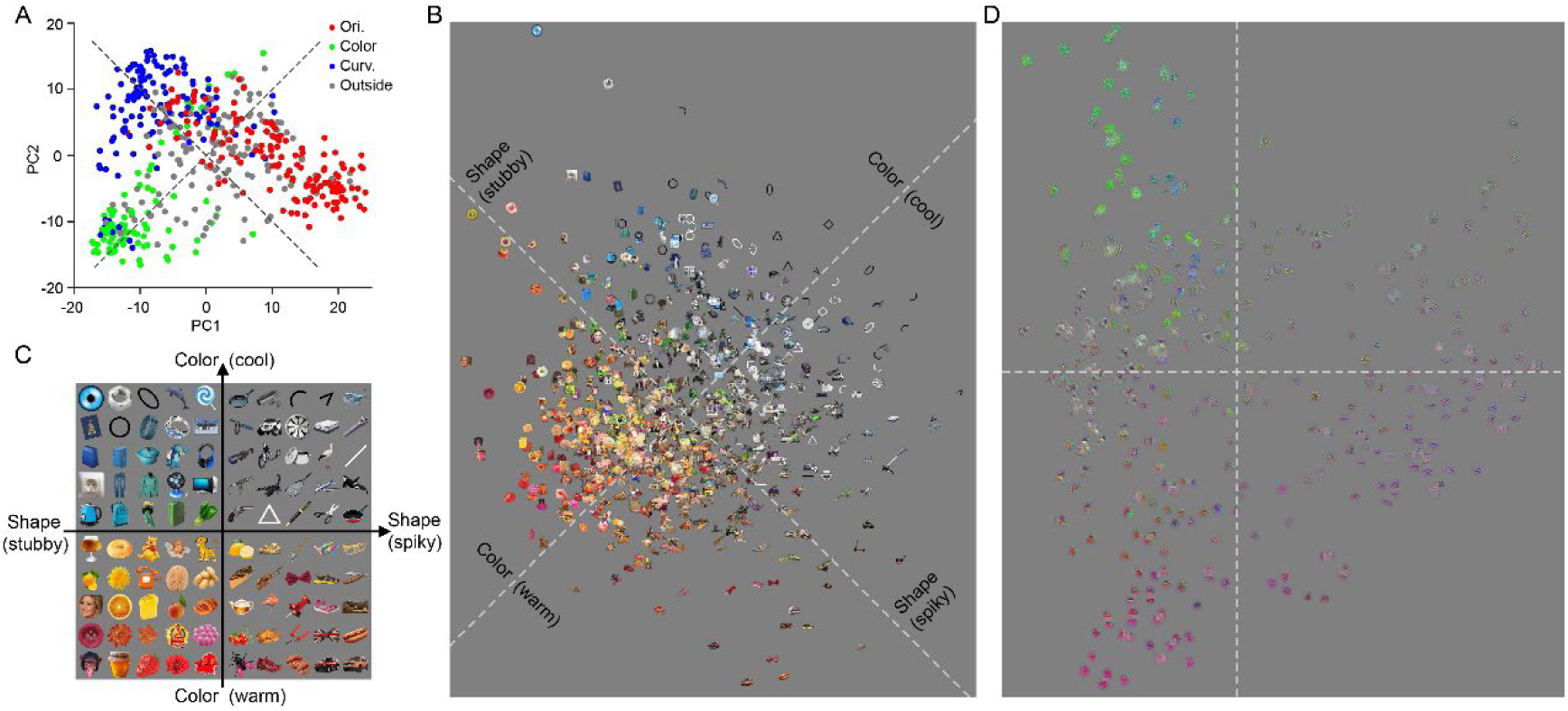
V4 population responses in stimulus space. (for Figure 4) **A.** PCA was carried out for responses of V4 neurons’ (n=497) in stimulus dimension, including 800 images of natural objects and 50 images of geometric figures. Neurons were plotted in PC1-PC2 space. PC1 and PC2 each accounted for 16.6% and 9.1% variance in the responses. Similar to those shown in Figure 4C, neurons from different functional domains were clustered in different locations. Dotted lines are estimated dimensions for shape and color, which have a 45 deg angle with the PC axis. **B.** Stimulus images plotted in PC1-PC2 space according to their weight on these two axis. The rotated feature axes for color and shape are indicated as dotted lines. **C.** Representative images from B are replotted in the rotated features space (dotted lines in B) for better illustration. **D.** Similar to B, but for SI images plotted according to their weight on PCs shown in Figure 4C.

**Figure S5.**
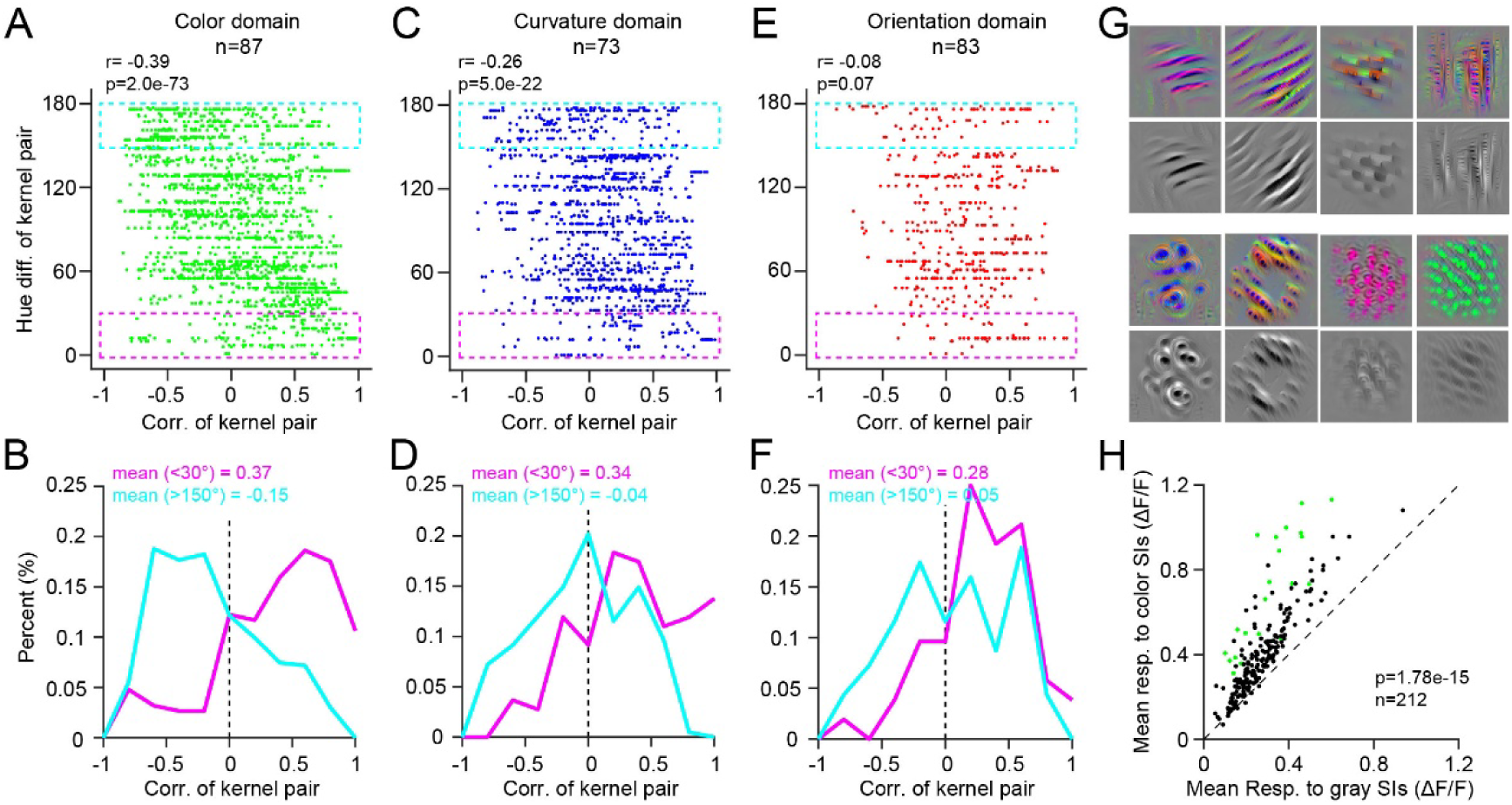
V4 neurons’ color inputs (for Figure 6) Similar to the analysis shown in Figure 6, we estimated how color kernels in L1 contributed to the modeled V4 color neurons. For each color neuron, we calculated the contribution vector when 800 stimulus images were input. **A.** For neurons in color domain, the color difference between L1 color kernels (Y-axis) is negatively correlated with the similarity of their contributions to the cell’s response (X-axis) (r=-0.39). The dashed box is quantified in B. **B.** For L1 color kernel pairs with similar color, their contributions to color cells’ responses are positively correlated (pink curve, from pink box in A). For L1 color kernel pairs with large color differences, their contributions to color cells’ responses are negatively correlated (cyan curve, from cyan box in A). **C&D**. Similar to A&B, but for neurons in curvature domains. Color differences of kernels are negatively correlated with their contributions to curvature neurons (r=-0.26). **E&F**. Similar to A&B, but for neurons in orientation domains. The correlation was insignificant. **G.** Color SIs and gray SIs for 4 orientation neurons (top), 2 curvature neurons and 2 color neurons (bottom). The gray SI was obtained using the same procedure for SI, but force the RGB values equal. It can be seen that gray SIs have the similar pattern as standard SIs. **H**. Neurons’ response to colored SI and gray SIs are strongly correlated. Not only color neurons (green dots) but also orientation and curvature neurons (black dots) have stronger responses to color SIs than gray SIs. Each response is an average of responses to all SI images.

## References

Bashivan P, Kar K, and DiCarlo J. (2019) Neural population control via deep image synthesis. Science, 364(6439)

Burg MF et al. (2024) Most discriminative stimuli for functional cell type clustering. arXiv: 2401.05340v2.

Bushnell BN & Pasupathy A (2012) Shape encoding consistency across colors in primate V4. Journal of Neurophysiology 108:1299–1308.

Conway BR, Moeller S, Tsao DY (2007) Specialized color modules in macaque extrastriate cortex. Neuron 56:560–573.

Desimone R, Schein SJ. (1987) Visual properties of neurons in area V4 of the macaque: sensitivity to stimulus form. Journal of Neurophysiology 57:835–868.

De Yoe EA, Felleman DJ, Van Essen DC, McClendon E (1994) Multiple processing streams in occipitotemporal visual cortex. Nature 371:151–154.

Fang C, Yan K, Liang C, Wang J, Cai X, Zhang R, Lu HD. (2022) Function-specific projections from V2 to V4 in macaques. Brain Struct Funct 227: 1317–1330.

Felleman DJ, Xiao Y, McClendon E (1997) Modular organization of occipito-temporal pathways: cortical connections between visual area 4 and visual area 2 and posterior inferotemporal ventral area in macaque monkeys. J Neurosci 17:3185–3200.

Franken TP, Reynolds JH (2021) Columnar processing of border ownership in primate visual cortex. Elife 10:e72573.

Friedman HS, Zhou H, and von der Heydt R (2003) The coding of uniform colour figures in monkey visual cortex. J Physiol, 548.2:593–613.

Garg AK, Li P, Rashid MS, Callaway EM (2019) Color and orientation are jointly coded and spatially organized in primate primary visual cortex. Science 364:1275–1279.

Ghose GM, Ts’o DY. (1997) Form processing modules in primate area V4. Journal of Neurophysiology 77:2191–2196.

Hatanaka G, Inagaki M, Takeuchi RF, Nishimoto S, Ikezoe K, Fujita I. (2022) Processing of visual statistics of naturalistic videos in macaque visual areas V1 and V4. Brain Structure and Function 227:1385–1403.

Hu JM, Song XM, Wang Q, Roe AW (2020) Curvature domains in V4 of macaque monkey. Elife 9:e57261.

Jiang R, Andolina IM, Li M, Tang S (2021) Clustered functional domains for curves and corners in cortical area V4. eLife 10:e63798.

Kim T, Kamath R, Hatanaka G, Namima T, Dylla C, Bair W, and Pasupathy A (2025) Functional clusters for shape, texture, and motion encoding in macaque V2. J Neurosci, 46(14). 10.1523/jneurosci.1994-25.2026

Kotake Y, Morimoto H, Okazaki Y, Fujita I, and Tamura H. (2009) Organization of Color-Selective Neurons in Macaque Visual Area V4. Journal of Neurophysiology 102:15–27.

Li, M., Liu, F., Juusola, M., and Tang, S. (2014). Perceptual color map in macaque visual area V4. J. Neurosci. 34, 202–217.

Li P, Zhu S, Chen M et al (2013) A Motion Direction Preference Map in Monkey V4. Neuron 78:376–388.

Lu L, Yin J, Chen Z, Gong H, Liu Y, Qian L, Li X, Liu R, Andolina IM, Wang W. (2018) Revealing Detail along the Visual Hierarchy: Neural Clustering Preserves Acuity from V1 to V4. Neuron 98:417–428.

Namima T, Kempkes E, Zamarashkina P, Owen N, Pasupathy A. (2025) High-Density Recording Reveals Sparse Clusters (But Not Columns) for Shape and Texture Encoding in Macaque V4. The Journal of Neuroscience 45 (5) e1893232024.

McClurkin JW & Optican LM. (1996) Primate striate and prestriate cortical neurons during discrimination. I. simultaneous temporal encoding of information about color and pattern. Journal of Neurophysiology 75:481–495.

Nakamura H, Gattass R, Desimone R, Ungerleider LG (1993) The modular organization of projections from areas V1 and V2 to areas V4 and TEO in macaques. J Neurosci 12:3681–3691.

Nigam S, Pojoga S, Dragoi V (2021) A distinct population of heterogeneously color-tuned neurons in macaque visual cortex. Science Advances 7:eabc5837.

Pasupathy A, Popovkina D, Kim T. (2020) Visual Functions of Primate Area V4. Annu. Rev. Vis. Sci. 6:20.1–20.23.

Roe AW, Chelazzi L, Connor CE, Conway BR, Fujita I, Gallant JL, Lu H, Vanduffel W (2012) Toward a unified theory of visual area V4. Neuron 74:12–29.

Schein SJ, Marrocco RT, de Monasterio FM. (1982) Is there a high concentration of color-selective cells in area V4 of monkey visual cortex? J Neurophysiol 47: 193–213.

Srinath R, Emonds A, Wang Q et al (2021) Early emergence of solid shape coding in natural and deep network vision. Curr Biol 31:51–65.e5.

Tanaka M, Weber H, Creutzfeldt OD. (1986) Visual properties and spatial distribution of neurons in the visual association area on the prelunate gyrus of the awake monkey. Exp Brain Res 65: 11–37.

Tang R, Song Q, Li Y et al (2020) Curvature-processing domains in primate V4. Elife 9:e57502.

Tanigawa H, Lu HD, Roe AW (2010) Functional organization for color and orientation in macaque V4. Nat Neurosci 13:1542–1548.

Vanduffel W, Tootell RBH, Schoups AA, Orban GA (2002) The organization of orientation selectivity throughout macaque visual cortex. Cerebral Cortex 12:647–662.

Walker, et al. (2019) Inception loops discover what excites neurons most using deep predictive models. Nature Neuroscience 22:2060–2065

Wang, H., Huang, L., Du, C., Li, D., Wang, B., & He, H. (2020). Neural encoding for human visual cortex with deep neural networks learning “what” and “where”. IEEE Transactions on Cognitive and Developmental Systems, 13(4), 827–840.

Wang T, Lee TS, Yao H, Hong J, Li Y, Jiang H, Andolina IM & Tang S. (2024) Large-scale calcium imaging reveals a systematic V4 map for encoding natural scenes. Nature Communications 15:6401.

Wang J, Zhang R, Cai X, Tang R, Dai Z, Lu HD (2025) Stimulus-driven rivalry among V1 neurons. Progress in Neurobiology. 254:102820, doi: 10.1016/j.pneurobio.2025.102820

Willeke KF, et al. (2026) Deep learning-driven characterization of single cell tuning in primate visual area V4 supports topological organization. eLife 15 :RP109875. 10.7554/eLife.109875.1

Xiao Y, Zych A, Felleman DJ (1999) Segregation and convergence of functionally defined V2 Thin stripe and interstripe compartment projections to area V4 of macaques. Cereb Cortex 9:792–804.

Zhang R, Wang J, Cai X, Tang R, Lu HD. (2025) Dynamic grouping of ongoing activity in V1 hypercolumns. Neuroimage. 310:121157. Doi: 10.1016/j.neuroimage.2025.121157.

Zhang Y, Schriver KE, Hu JM, Roe AW (2023) Spatial frequency representation in V2 and V4 of macaque monkey. eLife 12:e81794.

Zeki S (1983) The distribution of wavelength and orientation selective cells in different areas of monkey visual cortex. Proc R Soc Lond B Biol Sci 217:449–470.

Zeki S, Shipp S (1989) Modular connections between areas V2 and V4 of macaque monkey visual cortex. Eur J Neurosci 1:494–506.

Bashivan, P., Kar, K., & DiCarlo, J. J. (2019). Neural population control via deep image synthesis. Science, 364(6439), eaav9436.

Klindt, D., Ecker, A. S., Euler, T., & Bethge, M. (2017). Neural system identification for large populations separating “what” and “where”. Advances in Neural Information Processing Systems, 30.

Li, P., Hastie, T. J., & Church, K. W. (2006, August). Very sparse random projections. In Proceedings of the 12th ACM SIGKDD international conference on Knowledge discovery and data mining (pp. 287–296).

Güçlü, U., & van Gerven, M. A. (2015). Deep neural networks reveal a gradient in the complexity of neural representations across the ventral stream. Journal of Neuroscience, 35(27), 10005–10014.

